# SpeedSeq: Ultra-fast personal genome analysis and interpretation

**DOI:** 10.1101/012179

**Authors:** Colby Chiang, Ryan M Layer, Gregory G Faust, Michael R Lindberg, David B Rose, Erik P Garrison, Gabor T Marth, Aaron R Quinlan, Ira M Hall

## Abstract

Comprehensive interpretation of human genome sequencing data is a challenging bioinformatic problem that typically requires weeks of analysis, with extensive hands-on expert involvement. This informatics bottleneck inflates genome sequencing costs, poses a computational burden for large-scale projects, and impedes the adoption of time-critical clinical applications such as personalized cancer profiling and newborn disease diagnosis, where the actionable timeframe can measure in hours or days. We developed SpeedSeq, an open-source genome analysis platform that vastly reduces computing time. SpeedSeq accomplishes read alignment, duplicate removal, variant detection and functional annotation of a 50X human genome in <24 hours, even using one low-cost server. SpeedSeq offers competitive or superior performance to current methods for detecting germline and somatic single nucleotide variants (SNVs), indels, and structural variants (SVs) and includes novel functionality for SV genotyping, SV annotation, fusion gene detection, and rapid identification of actionable mutations. SpeedSeq will help bring timely genome analysis into the clinical realm.

**Availability:** SpeedSeq is available at https://github.com/cc2qe/speedseq.

## Background

Technical advances in second-generation DNA sequencing technologies have reduced both the cost and time required to generate whole-genome sequencing (WGS) data for clinical and research applications. At the time of writing, a human genome can be sequenced to ∼30X coverage in ∼3 days for ∼$1000. These improvements afford new opportunities in healthcare and academic research to survey the human genome with unprecedented depth and scope. Personalized clinical WGS can reveal causative mutations in patients who present with rare idiopathic congenital disorders, for whom performing dozens of individual genetic assays is financially impractical. It can also elucidate the unique combination of genetic aberrations in a patient’s tumor, informing clinical decisions when selecting from a growing number of targeted pharmaceutical agents designed to specifically neutralize oncogenic drivers. A growing list of success stories as well as increasing readiness for insurance reimbursement have made WGS an attractive diagnostic tool for clinicians and patients alike[1, 2, 3, 4, 5].

However, major barriers in the speed and reproducibility of WGS data interpretation have hindered widespread adoption of WGS in clinical and research applications. Using a standard pipeline based on BWA[6], GATK[7], SAMtools[8] and Picard[9], computational processing of a 50X human genome from raw sequence data to variant calls on a single 16-core server requires 68-94 hours, depending on software versions and the level of parallelization. After variant detection, variants must be annotated and interpreted in the context of an extremely large and diverse body of relevant information including genomic features, known disease-causing alleles, gene ontologies, various genotype/phenotype databases, and functional variant prediction software. Due to the complexities and ambiguities inherent to this process, current pipelines for variant interpretation necessitate extensive hands-on involvement from informatics specialists and genetic counselors to distinguish pathogenic from benign mutations, and to prioritize candidates. This labor-intensive and time-consuming process is estimated to require up to 100 hours of manual curation per patient[10]. The time required to go from raw data to diagnosis is a crucial issue because delays can significantly worsen patient outcome, particularly for newborn diagnosis where many treatable metabolic disorders have a brief therapeutic window. In previous studies, this informatics bottleneck has been circumvented in one of three ways: by surveying a small subset of genetic variants previously implicated in certain diseases[11], by performing low-coverage surveys of structural variation (SV)[12], or by allocating nearly two months for detailed analysis[13].

Besides clinical considerations, WGS data analysis time is an increasingly important factor for research projects. Large-scale WGS-based projects involving thousands of individuals are becoming common as DNA sequencing throughput increases and costs drop, and such studies demand a rapid and reproducible genome analysis workflow to achieve high quality results within time constraints and computational resource limitations. For smaller projects and isolated experiments, an intuitive and rigorously validated pipeline can streamline routine analysis tasks such as alignment and variant calling, liberating resources to address project-specific research aims.

With these objectives in mind, we present SpeedSeq, a comprehensive and intuitive analysis pipeline for characterizing and interpreting genetic variation in less than 24 hours per genome on a single low-cost commodity server.

## Results and discussion

### Software overview

SpeedSeq is a modular suite of open-source software designed for rapid whole-genome variant detection and interpretation under a variety of experimental paradigms including individual, family, tumor/normal or population scale sequencing (Fig. 1). SpeedSeq achieves superior processing efficiency without compromising performance by engineering non-dependent pipeline components to run simultaneously, by multi-threading external applications, and by empirically partitioning the genome for balanced parallelization. Its core components - `speedseq align`, `speedseq var/somatic`, and `speedseq sv` - can be run independently or sequentially. Internally, SpeedSeq uses BWA-MEM[14] for read alignment, SAMBLASTER[15] for in-memory duplicate marking, and Sambamba[16] for multi-threaded sorting and indexing to produce BAM alignment files for downstream analyses. SpeedSeq runs a parallelized implementation of FreeBayes[17] to detect single nucleotide variants (SNVs) and small insertion/deletion variants (indels), with optimized parameters for either germline or somatic mutation detection. SpeedSeq detects genome structural variation (SV) using a combination of read-pair and split-read data using LUMPY[18] and read-depth analysis using CNVnator[19], and includes a novel SV breakpoint genotyper (SVTyper). SpeedSeq produces VCF files that are annotated by the Ensembl Variant Effect Predictor (VEP)[20] and can be loaded into the GEMINI genome interpretation software[21] for interactive exploration or automated querying. We added new features to GEMINI that enable proper SV annotation and interpretation of variants in cancer samples.

**Figure 1.**
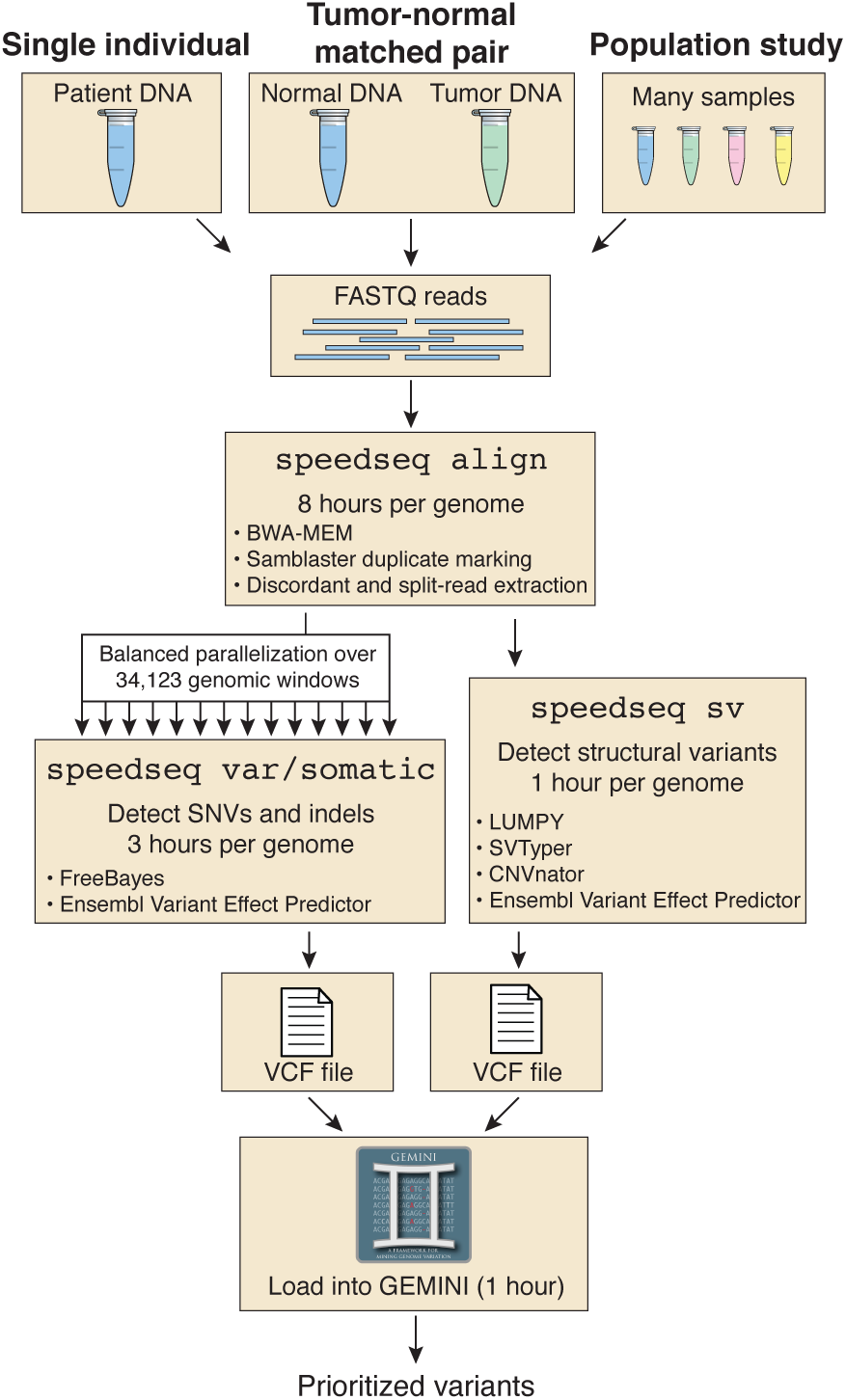
SpeedSeq workflow. SpeedSeq can convert raw, FASTQ reads from nearly any experimental design into prioritized variants in ∼13 hours for a 50X human dataset. Its modular architecture consists of “speedseq align” to generate BAM files, “speedseq var” to detect SNVs and indels, and “speedseq sv” to detect structural variants. The function “speedseq somatic” uses specialized logic to detect somatic SNVs and indels in a tumor-normal pair. VCF files may then be loaded into the GEMINI database framework to produce prioritized variant calls. Timings shown in this diagram were performed the 50X NA12878 human dataset from the Illumina Platinum Genomes project (European Nucleotide Archives; ERP001960) on a single machine with 128 GB RAM and two Intel Xeon E5-2670 processors.

Many of SpeedSeq’s core algorithms have been published previously and are available as standalone tools; however, several key aspects of this work are new. First, we have devised a method to perform read-alignment, duplicate marking and coordinate sorting in one data processing step without landing intermediate files, which simplifies workflows and greatly reduces computing time and disk requirements. Second, we have developed a Bayesian SV breakpoint genotyping algorithm (SVtyper) that produces accurate genotypes for germline breakpoints and quantitative variant allele frequency estimates for somatic breakpoints, information that is crucial for clinical SV interpretation yet not provided by most SV detection tools. Third, we have developed a new algorithm for detecting and interpreting cancer gene fusions from genome sequencing data. Fourth, we have developed a framework for highly parallelized SNV and indel detection using FreeBayes, which is based on independent processing of 34,123 variably-sized genomic intervals selected to exhibit similar read depth levels. Fifth, to test our software on real WGS data, we have developed a novel transmission-based approach for measuring algorithm performance using large human pedigrees – in this case the 17 member CEPH 1463 pedigree that includes the well-studied NA12878 individual. Finally, we have made notable improvements to existing software. SpeedSeq implements a streamlined LUMPY workflow that achieves a 32-fold speedup over the published version[18], a custom-designed parallelized build of CNVnator that is ∼31-fold faster than the original algorithm, and a significantly improved version of GEMINI that is designed for clinical interpretation of both germline and somatic variants – including structural variants and fusion genes – and that interacts with cancer drug databases to facilitate identification and interpretation of clinically actionable mutations.

### Software runtimes

To test SpeedSeq’s performance, we utilized the Illumina Platinum Genomes data (European Nucleotide Archive: ERP001960), which comprise ∼50X WGS datasets for each of the 17 members of the three generation CEPH 1463 pedigree. For single-genome analyses, we focus on the well-studied NA12878 genome, which is the mother in this pedigree. To facilitate reproduction of these performance tests and to enable further testing and optimization by the community, we have made SpeedSeq available as a public Amazon Machine Instance, which can be cloned for multiplex analysis of numerous datasets. SpeedSeq translates raw sequence data into prioritized, clinically interpretable SNV, indel, and SV calls in ∼13 hours for NA12878 (Fig. 1) using default software parameters and a single 16-core server (allowing 32 threads) with 128 GB of RAM (current cost: <$10,000). This represents at minimum a several-fold speed increase over current practices, reducing computing requirements to fit within clinically accessible time constraints. In contrast to cloud-based software applications that achieve processing speed by harnessing a distributed network of independent CPUs, SpeedSeq is designed to process data on a single multi-core server. This model offers the benefit of local data processing, which is an attractive option for applications within the healthcare industry, where patient privacy and data security are paramount. However, the general architecture of SpeedSeq is inherently scalable, allowing flexibility for cloud implementation of most modules, and we expect that substantial speed increases can be achieved through distributed computing. We further note that in the context of a single WGS dataset, increased parallelization yields diminishing returns because runtimes are bounded by difficult-to-parallelize steps such as BAM file sorting and duplicate marking (Fig. 1).

SpeedSeq’s fast runtimes are achieved through several parallelization and synchronization steps. A considerable reduction in processing time is achieved through duplicate marking with our SAMBLASTER program[15], which operates on streamed input from the aligner and can harvest idle CPU cycles that are periodically released by BWA, decreasing duplicate marking time to ∼12 minutes (a process that takes 8.0 hours using Picard). SpeedSeq also uses Sambamba for multi-threaded BAM manipulation, providing a 1.6-fold speed increase for sorting and compression relative to the widely-used SAMtools package.

Parallelized SNV and indel calling is achieved by running FreeBayes on 34,123 variably-sized genomic windows that have been selected to contain similar numbers of reads based on aggregate aligned read depth of the 17 CEPH individuals used in this study. Our binning of the human genome by empirical depth accomplished two objectives relevant to improving performance. First, we identified 15.6 Mb (0.6%) of the GRCh37 genome build with consistently excessive read depth, presumably due to paralogous genomic segments absent from the reference assembly. By excluding these regions we avoided calling variants in areas where reference artifacts violate assumptions of ploidy and mappability, while simultaneously reducing computational time by ignoring massive data pileups. Secondly, the depth-based exclusion strategy allowed us to construct variable-width bins to balance parallelized FreeBayes variant calling (Supplementary Fig. 1). While this static set of excluded regions was based on a single large family, the depth cutoff accommodated a 2-fold increase in copy number relative to the reference genome - corresponding to a homozygous duplication (4 copies) in a genomic region that is single copy in the reference - minimizing the bias toward fixed polymorphism in that family. In the future, we envision that excluded genomic regions will be defined by a much larger set of high-quality human genomes from diverse populations.

### Germline SNVs and indels

The central goal of SpeedSeq is to achieve fast runtimes without sacrificing variant detection sensitivity or accuracy. We therefore set out to rigorously test variant detection performance under typical usage scenarios. Perhaps the most common and straightforward WGS analysis is to map germline SNVs and indels. If we accept as truth the 2,803,144 SNV and 364,031 indel calls reported for NA12878 by the Genome in a Bottle (GIAB), SpeedSeq achieves a sensitivity of 99.9% and 89.9% for SNVs and indels, respectively, and has acceptably low FDRs (0.4% and 1.1%) (Fig. 2a,b). These detection rates exceed the GATK Unified Genotyper’s (GATK-UG) sensitivity (99.7%, 89.0%; SNVs, indels) with a similar FDR (0.5%, 1.0%; SNVs, indels). The GATK Haplotype Caller (GATK-HC) shows superior indel detection sensitivity (99.8%, 95.7%; SNVs, indels) with lower FDR for both variant types (0.2%, 0.6%; SNVs, indels). SpeedSeq’s implementation of FreeBayes therefore exhibits comparable – albeit slightly inferior – performance relative to GATK-HC when tested on the GIAB callset. However, since the GIAB truth set was primarily derived from GATK-based analyses, it is likely to be biased towards GATK tools. We therefore assessed SpeedSeq’s performance against a second truth set derived from the Illumina Omni 2.5 SNP array results for NA12878, which contains 2,177,040 informative markers that are expected to be unbiased relative to the different tools (albeit enriched for easy-to-genotype markers). All variant callers show similar performance against the Omni microarray truth set, with SpeedSeq attaining the highest sensitivity at the minor expense of specificity compared to GATK-UG or GATK-HC (Fig. 2c). These results show that SpeedSeq detects germline SNVs and indels with competitive sensitivity and specificity to GATK, a widely-used software package that is generally considered to be the industry standard.

**Figure 2.**
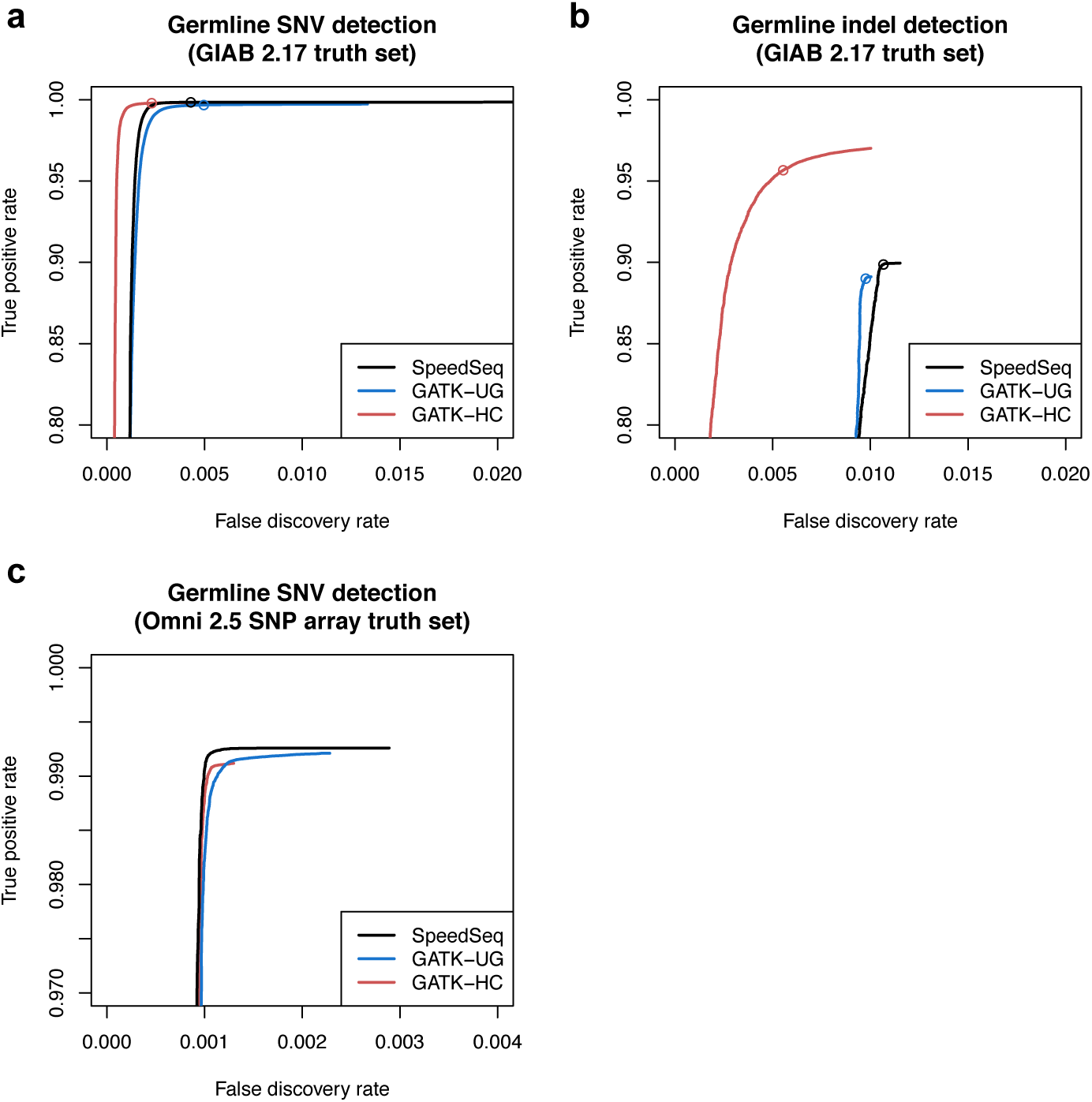
Germline SNV and indel detection performance. Receiver operating characteristic (ROC) curves comparing the performance of SpeedSeq (black lines) to GATK Haplotype Caller (red lines) and GATK Unified Genotyper (blue lines) in detecting germline SNVs and indels in the 50X NA12878 human dataset, with quality score as the varying parameter. (a) SNVs and (b) indels reported by each algorithm were benchmarked against the NA12878 callset from the Genome in a Bottle Consortium (version 2.17), with open circles denoting the quality thresholds for sensitivity and specificity metrics reported in the main text (see Germline SNVs and Indels). (c) SNVs detected by each algorithm benchmarked against an Omni 2.5 SNP array (note the zoomed axes scales in comparison to panels a and b).

### Somatic SNVs and indels

Profiling somatic mutations in cancer samples is one of the most common clinical applications of WGS, and is a clear example of a situation where fast analysis turnaround time has the potential to directly affect patient care. Somatic mutation detection is a difficult bioinformatic problem that requires specialized logic to weigh evidence from both samples in a tumor-normal matched pair, since classifying a variant as somatic requires evidence not only that the tumor carries the variant, but also that the normal sample does not. Moreover, many tumors exhibit subclonal heterogeneity and/or chromosomal aneuploidy, both of which violate the assumptions of diploidy made by most variant callers. Finally, clinically relevant cancer WGS requires reliable detection of variants at or below 10% variant allele frequency (VAF), since actionable mutations may reside in minor subclones. To overcome these challenges, we tuned FreeBayes’s algorithmic parameters and filtering criteria to jointly analyze a tumor-normal pair, probing for subclonal variants as low as 5% VAF in the tumor with no assumption of diploidy. To filter out artifacts and misclassified variants, we established effective filtering criteria for SpeedSeq, requiring variants to meet a log odds threshold for both the tumor and normal samples based upon FreeBayes genotype likelihood scores.

We next sought to assess SpeedSeq’s performance. It is currently infeasible to conduct a realistic comparison of somatic mutation detection performance using bona fide cancer WGS data and an adequately large truth set, because comprehensive and unbiased somatic mutation callsets do not exist. For this reason, most cancer WGS tool comparisons have relied either on entirely simulated data or artificial injection of mutations into real data, both of which have limited value. We instead devised a novel strategy for assessing somatic mutation detection on real WGS data using the CEPH 1463 pedigree datasets. To emulate a WGS dataset from a heterogeneous tumor sample with somatic mutations present at varying allele frequencies, we pooled raw data from each of the 11 grandchildren in equal proportions to generate a single 50X chimeric sample. For performance tests, we define this chimeric dataset as the tumor sample and the father’s dataset (NA12877) as the normal sample, and we define “somatic mutations” as variants that are present in the mother (NA12878) and absent from the father. Assuming Mendelian segregation, somatic mutations that are heterozygous in the mother are expected to be present in the chimeric dataset at a range of VAFs defined by a binomial distribution centered at 0.25, and variants that are homozygous in NA12878 will be present at a VAF of 0.5. This experimental design provided us with a somatic truth set of 875,206 variants with known VAFs ranging from 0.05 to 0.5 matching the expected distribution (Fig. 3a).

**Figure 3.**
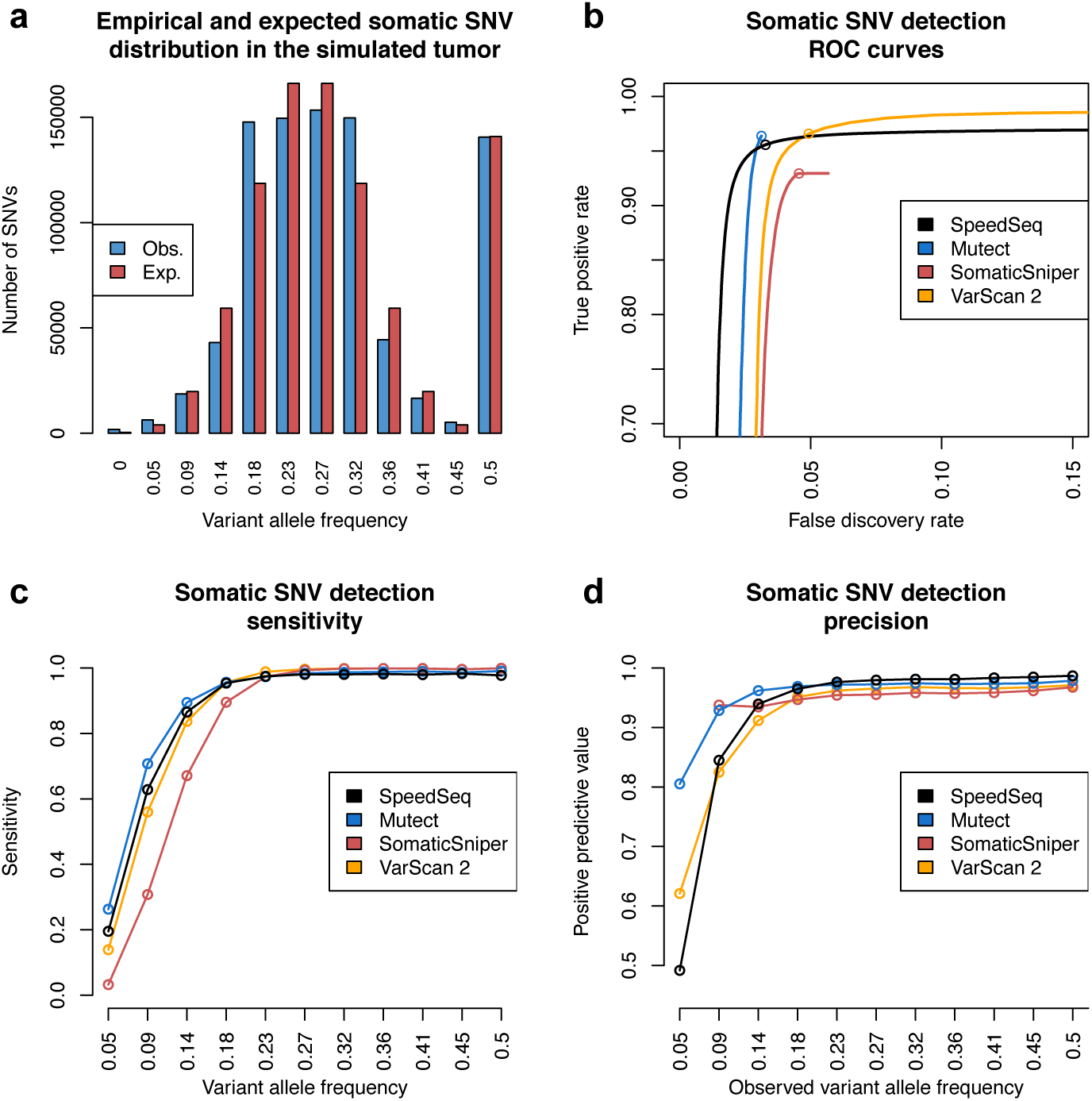
Somatic SNV detection performance of low frequency variants in a simulated tumor-normal pair. (a) Somatic variants in the simulated 50X tumor dataset (a mixture of 11 grandchildren from the CEPH 1463 pedigree) exhibit a range of variant allele frequencies in accordance with the expected binomial distribution. (b) Receiver operating characteristic (ROC) curves comparing the performance of SpeedSeq (black line) to Mutect (blue line), SomaticSniper (red line), and VarScan 2 (orange line). Open circles denote quality thresholds used for sensitivity and precision plots showing SNV detection sensitivity (c) and precision (d) over the range of variant allele frequencies in the simulated tumor.

Using this evaluation paradigm, we compared SpeedSeq’s performance to three other leading somatic variant calling tools: MuTect[22], SomaticSniper[23], and VarScan 2[24]. SpeedSeq recalled 96.6% of the somatic variants in the chimeric tumor with a FDR of 3.3%, outperforming SomaticSniper in both sensitivity and specificity, and delivering competitive performance against MuTect and VarScan 2 (Fig. 3b). For variants with a VAF of less than 0.2, SpeedSeq was more sensitive than both VarScan 2 and Somatic-Sniper, detecting SNVs at 0.09 VAF with a sensitivity of 62.8% and a positive predictive value of 84.5% (Fig. 3c,d). These results demonstrate that SpeedSeq identifies somatic SNVs and indels with similar performance as the current best-in-class mutation callers.

### Strategy for detecting genome structural variation

Comprehensive ascertainment of structural variation (SV) is a critical yet technically diffcult component of diagnostic sequencing. SV is a broad class of genome variation that includes copy number variants (CNVs) such as deletions and duplications, balanced rearrangements (e.g., inversions or translocations), complex rearrangements, and mobile element insertions. Although SVs are rare in number compared to SNVs and indels – a typical human genome harbors approximately 5,000 germline SVs that are detectable with current technologies – they can affect large genomic segments and can have severe functional consequences due to their ability to disrupt genes, cause gene fusions, rearrange regulatory elements, and alter gene dosage.

SV poses two key challenges for clinical implementation of WGS. First, for various technical reasons, SV is recognized to be by far the most difficult form of variation to detect reliably[25]. Second, functional interpretation of SV requires specialized logic due to the variable size and diverse configurations that SVs exhibit, and because SV breakpoints are often mapped imprecisely. Due to these challenges, few established genome analysis pipelines attempt to rigorously detect and interpret SV. Yet, given the large body of literature supporting a role for SV in various human diseases, perhaps most notably in the etiology of genomic disorders and cancer, it is clear that sensitive and accurate SV detection is a prerequisite for clinical WGS.

SpeedSeq uses LUMPY as the core component of its structural variation module. LUMPY identifies deletions, duplications, inversions and other rearrangements by combining multiple evidence types (e.g. read-pair and split-read) within a probabilistic framework, and has recently been demonstrated to match or outperform other leading tools for detecting germline and somatic SVs across diverse variant types and allele frequencies[18]. SpeedSeq includes significant improvements to the LUMPY workflow to improve runtimes and usability. SpeedSeq simplifies LUMPY execution by automatically pre-processing the BAM files for insert size distributions and configuring multiple samples and libraries for joint variant calling. Extraction of discordant read-pair and split-read alignments by SAMBLASTER during duplicate marking enables extremely fast LUMPY runtimes, requiring merely ∼74 minutes to call SV breakpoints on the 50X NA12878 dataset. This is in contrast to 23.9 hours and 18.3 hours of processing time for two other leading SV detection tools, DELLY[26] and GASVPRO[27]. Finally, we have enhanced LUMPY’s output format to ease human readability and facilitate downstream analyses by producing an annotated VCF file as well as a standard BEDPE[28] file.

Besides mere detection, differentiating SV genotypes (SV) is a critical yet technically difficult component of and estimating breakpoint allele frequencies is crucial for assessing pathogenicity. For example, at haplosufficient loci, homozygous, but not heterozygous loss of function mutations can result in a pathogenic phenotype. At somatic variants, quantitative estimates of breakpoint allele frequencies allow inference of the fraction of tumor cells that carry a particular variant, which can help inform treatment options. Due to technical challenges, most SV detection algorithms do not provide SV genotypes nor quantitative estimates of breakpoint allele frequency. We therefore developed SVTyper, a Bayesian likelihood algorithm to genotype SV breakpoints based on reference and alternate read observations at each junction. This strategy serves as a complementary approach to genotyping deletions by read-depth analysis[29], as it can interrogate copy neutral events such as inversions or translocations as well as deletions that lack an observable reduction in depth due to their small size or the presence of repetitive sequence. SVTyper assumes a random sampling of reads at each breakpoint, with the number of reference and alternate observations being approximately equal in a heterozygote, and skewing toward either type of evidence in a homozygote, using Bayesian classifiers to ascribe confidence scores to each variant call (see Methods).

Finally, certain SV breakpoints contained within large repeats may be difficult or impossible to map precisely using current sequencing technologies. Notable examples include the recurrent pathogenic mutations that underlie a number of human genomic disorders[30]. When such variants alter the copy number of relatively large genomic segments (e.g., >1 kb), they can be detected via read-depth analysis. We have therefore packaged SpeedSeq with a well-tested CNV detection algorithm, CNVnator[19], that allows the user to optionally perform read-depth analysis in conjunction with LUMPY and SVTyper. To achieve acceptably fast runtimes, we implemented an optimized and parallelized version of CNVnator that processes the 50X NA12878 genome in ∼26 minutes compared to 13.5 hours for the original algorithm, representing a 31-fold speed increase (with virtually identical results). We also use CNVnator’s genotyping module to predict the copy number of the genomic interval affected by each LUMPY SV call, which aids functional interpretation.

### Structural variation performance benchmarking

Measuring SV detection performance on real data is difficult because there are not well-established truth sets that allow simple calculations of sensitivity and accuracy. If we accept the 1000 Genomes Project (1KGP) deletion callset for NA12878 as ground truth, SpeedSeq achieves a sensitivity of 61.9% (2,089/3,376) and positive predictive value of 60.8% (2,089/3,438) for detecting deletions, which is consistent with our recent comparative performance tests for LUMPY[18] and by inference shows that SpeedSeq achieves state-of-the-art SV detection relative to other tools. However, this test underestimates absolute performance because the 1KGP callset is known to contain false positives and false negatives. We therefore developed a multi-pronged validation strategy in which SVs reported by SpeedSeq in NA12878 could be validated either by overlap with split-read mapping of deep (∼30X) NA12878 long-read data (Fig. 4a, see Methods) or by overlap with deletions from the 1KGP pilot or phase 1 callsets (Fig. 4b). Based on this hybrid approach, SVs with quality scores of 100 or greater show a positive predictive value of 86.0% (2,823/3,282) (Fig. 4c,d). Virtually none of these SVs are likely to have been validated by random chance, as 100 permutations of the callset resulted in a validation rate of 0.073% (± 6.1E-3, 95% CI) (Fig. 4c). Moreover, these Phred-scaled quality scores, derived from SVTyper’s probabilistic framework, provide a tunable parameter through which users may refine callsets to a desired confidence threshold. By further requiring both paired-end and split-read support, users may generate an extremely high confidence callset of 1,663 SVs with a 97.8% validation rate.

**Figure 4.**
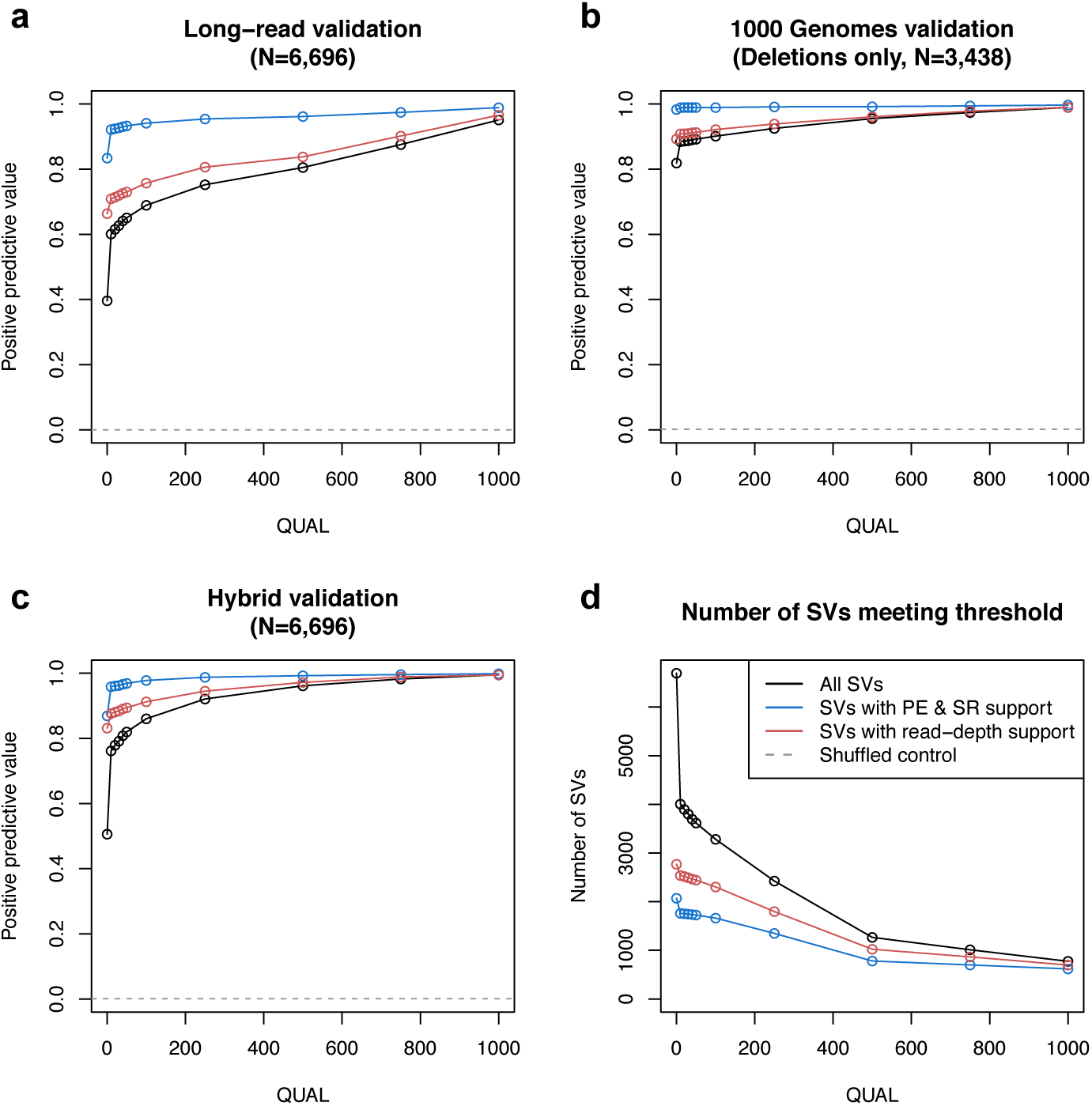
Structural variant validation by long-reads and 1000 Genomes Project data. SpeedSeq reported 6,696 structural variants (SVs) in the 50X NA12878 human dataset. The subsets of SVs with read-depth support from CNVnator (red lines) and with both paired-end and split-read support from LUMPY (blue lines) are displayed alongside the full set of reported variants (black lines) in each plot. (a) These variants were validated using deep (30X) long-read data from Pacific Biosciences or Illumina Moleculo technologies, showing positive predictive value for SVs at different quality thresholds and different evidence types. (b) We also validated the subset of 3,438 deletions reported by SpeedSeq against deletions reported in the pilot or phase 1 callsets of the 1000 Genomes Project. (c) We combined these two approaches into a hybrid validation metric, whereby SVs could be validated either by long-read support or by overlap with 1000 Genomes data. Gray hashed lines in panels a, b, and c denote the validation rate of 100 random permutations of the data. (d) The number of SVs meeting each quality thresholds.

As an independent measure of SV detection and genotyping performance, we developed a novel haplotype-based test that exploits the structure of the CEPH 1463 pedigree. We performed SNV-based phasing to produce haplotype lineage maps, allowing us to attribute an average of 63.0% of the mappable genome of each F_2_ individual to a particular founding grandparent (Fig. 5). We then used SpeedSeq to perform joint multi-sample SV analysis of the entire pedigree to identify SVs that could be unambiguously attributed to a grandparental haplotype. Joint calling alleviates ascertainment bias by allowing us to identify high confidence SVs that meet a minimum level of support in at least one of the 17 individuals, thus minimizing false positive SV calls, and to positively detect the presence of an SV in an individual based on a single discordant or split-read alignment, thus maximizing sensitivity for defining segregation patterns. This analysis resulted in 8,546 high confidence autosomal SV calls of which 1,722 could be assigned to a founding grandparental haplotype, with 8,397 predicted SV observations in the 11 grandchildren. SpeedSeq showed a detection sensitivity of 90.2% (7,578/8,397) for these SVs encompassing 1,660 of the 1,722 unique variants (Table 1).

**Figure 5.**
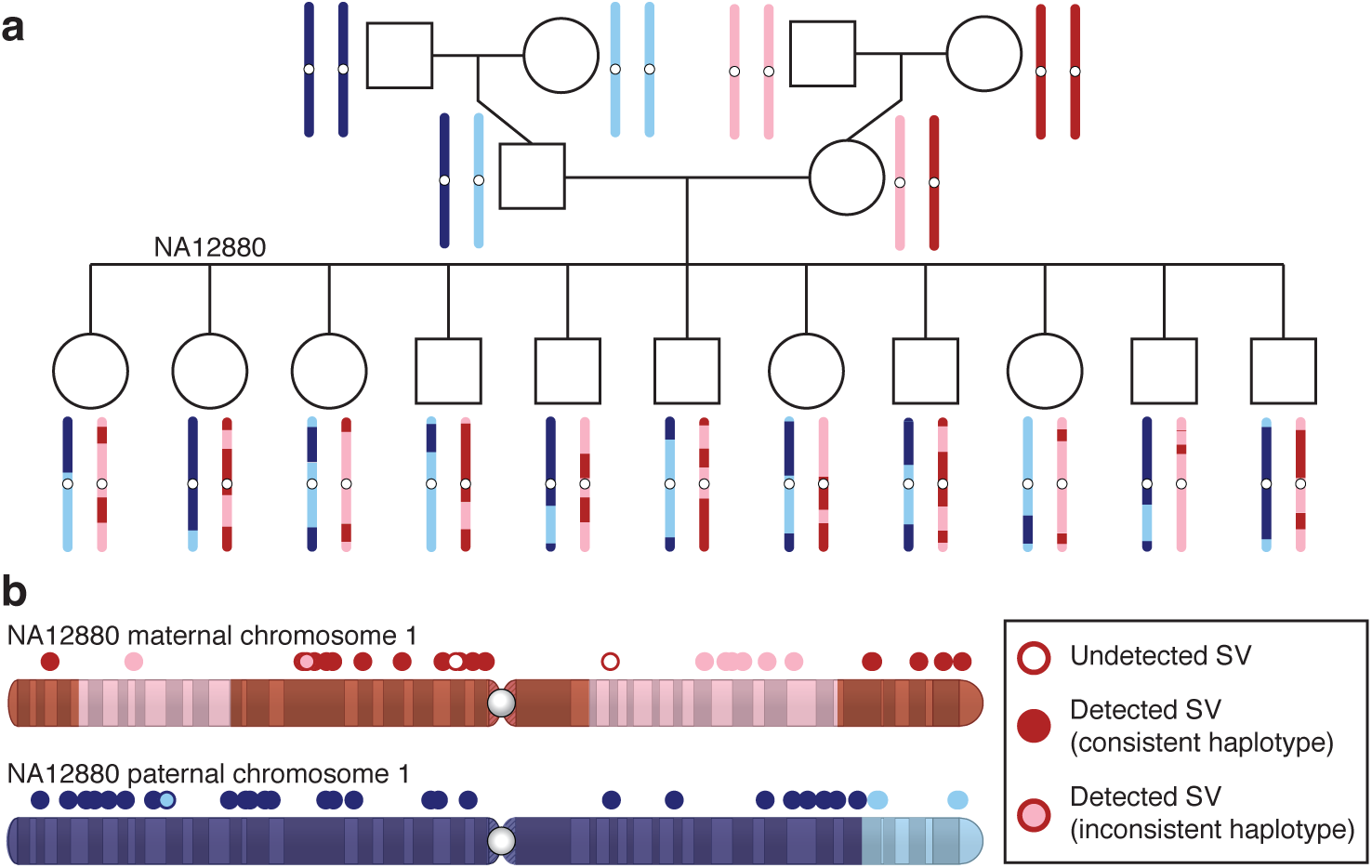
Haplotype-based structural variant validation. (a) The structure and haplotype maps of chromosome 1 for each member of the CEPH 1463 pedigree show the transmission of grandparental haplotype blocks to the grandchildren. (b) An example of this validation method on chromosome 1 of NA12880. The grandparental founder of each genome segment is denoted by the color of the chromosome (dark blue: paternal grandfather, light blue: paternal grandmother, pink: maternal grandfather, red: maternal grandmother). Circles above the chromosomes represent the SVs that are expected to be in NA12880 by transmission, with the color of the circumference denoting the founding grandparent predicted by the haplotype map, and the interior shading denoting the founding grandparent as defined by segregation of the SV. Hollow circles represent SVs that were predicted to be in NA12880 by the haplotype map, but undetected with SpeedSeq.

**Table 1.** Detection and genotyping accuracy of structural variants in the CEPH 1463 grandchildren. SNV-based haplotype phasing of genomic segments in the CEPH 1463 produced 1,722 variants (1,505 heterozygous and 217 homozygous) that were predicted in the 11 grandchildren. We used these 1,722 variants to assess detection sensitivity and genotyping accuracy across different variant classes and zygosities.

|  | Type | Correctly genotyped | % correctly genotyped | Detected | Detection sensitivity | Informative occurrences | Unique variants |
| --- | --- | --- | --- | --- | --- | --- | --- |
| All SVs | All | 7,201 | 95.0% | 7,578 | 90.2% | 8,397 | 1,722 |
|  | Deletion | 5,764 | 95.4% | 6,042 | 90.8% | 6,651 | 1,342 |
|  | Duplication | 859 | 93.7% | 917 | 89.5% | 1,025 | 217 |
|  | Inversion | 558 | 93.2% | 599 | 85.9% | 697 | 152 |
|  | Distant | 20 | 100.0% | 20 | 83.3% | 24 | 11 |
| Heterozygous SVs | All | 6,840 | 96.6% | 7,083 | 89.9% | 7,883 | 1,505 |
|  | Deletion | 5,418 | 96.0% | 5,641 | 90.4% | 6,240 | 1,173 |
|  | Duplication | 853 | 97.9% | 871 | 89.2% | 976 | 193 |
|  | Inversion | 549 | 99.6% | 551 | 85.7% | 643 | 128 |
|  | Distant | 20 | 100.0% | 20 | 83.3% | 24 | 11 |
| Homozygous SVs | All | 361 | 72.9% | 495 | 96.3% | 514 | 217 |
|  | Deletion | 346 | 86.3% | 401 | 97.6% | 411 | 169 |
|  | Duplication | 6 | 13.0% | 46 | 93.9% | 49 | 24 |
|  | Inversion | 9 | 18.8% | 48 | 88.9% | 54 | 24 |
|  | Distant | 0 | - | 0 | - | 0 | 0 |

To test whether the 1,722 informative SVs were representative of the dataset as a whole, and not of misleadingly high quality due to their ascertainment criteria, we assessed their validation rate as above using the 1KGP callset and long-read sequencing (Table 2). The 1,722 informative SVs have a similar validation rate as the remaining 6,734 SVs, which suggests that they are representative of overall callset quality.

**Table 2.**
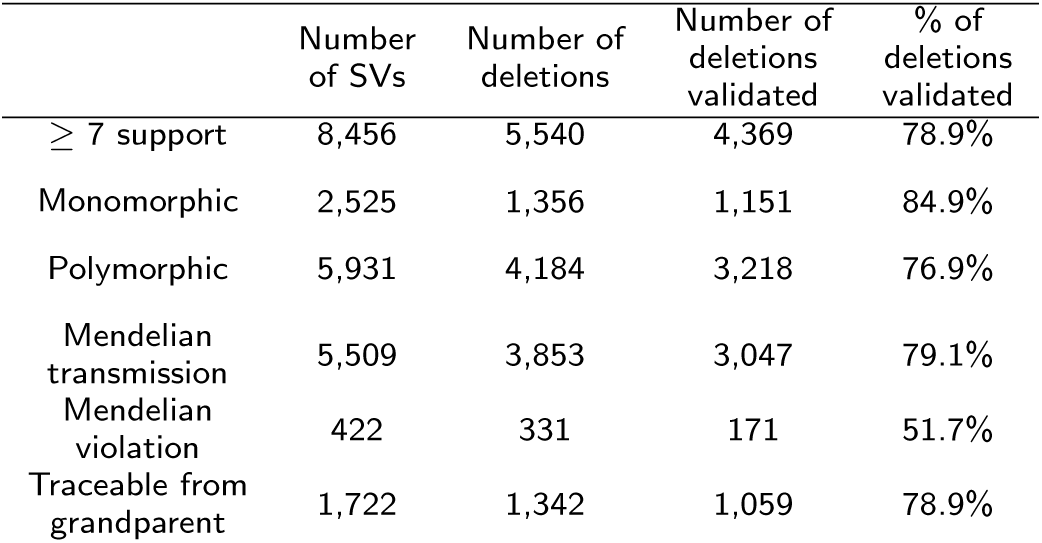
Filtering schematic for 8,456 high confidence SVs detected by SpeedSeq in the CEPH 1463 pedigree. Here we show the number of SVs and their validation efficiency at each stage of filtering to produce the 1,722 SVs that were used in SV benchmarking (Table 1).

|  | Number of SVs | Number of deletions | Number of deletions validated | % of deletions validated |
| --- | --- | --- | --- | --- |
| ≥ 7 support | 8,456 | 5,540 | 4,369 | 78.9% |
| Monomorphic | 2,525 | 1,356 | 1,151 | 84.9% |
| Polymorphic | 5,931 | 4,184 | 3,218 | 76.9% |
| Mendelian transmission | 5,509 | 3,853 | 3,047 | 79.1% |
| Mendelian violation | 422 | 331 | 171 | 51.7% |
| Traceable from grandparent | 1,722 | 1,342 | 1,059 | 78.9% |

To evaluate SV detection accuracy, we reasoned that true positive SV calls should exhibit presence/absence patterns within the pedigree that are consistent with familial relationships and haplotype transmission patterns, whereas false positive calls should exhibit random patterns. Based on this logic, two lines of evidence suggest that SpeedSeq’s SV detection accuracy is extremely high. First, only 422 of the 8,456 SVs (5.0%) have presence/absence patterns among the 17 individuals that violate the laws of Mendelian segregation. Of these, 51.7% are known variants based on 1KGP and/or long-read sequencing, indicating that false negatives are as common as false positives in causing Mendelian violations. Second, if we consider all observations of the 1,722 informative SVs among the 11 F_2_ individuals, only 3.1% contradict the SNV-based grandparental haplotype, and only 8.3% of variants show one or more contradictory observations. Since we expect false positive calls to co-segregate with grandparental haplotypes merely 50% of the time by chance, this result suggests that only a small fraction (521/8,397) of SV observations reported by SpeedSeq are false positives. To estimate SV detection sensitivity and genotyping accuracy, we constructed a SV truth set for F_2_ individuals using the observed haplotype transmission patterns. Based on inherited grandparental haplotypes, we derived the expected genotype for each of the 1,722 SVs (1,505 heterozygous and 217 homozygous) in each of the 11 F_2_ individuals, resulting in a truth set comprising 8,397 informative SV genotype observations. Of these, 7,833 are expected to be heterozygous for the alternate allele, 514 homozygous for the alternate allele. SVTyper reported the correct genotype 95.8% of the time for heterozygous variants and 70.1% for homozygous variants. This shows that SpeedSeq achieves clinically relevant levels of SV detection sensitivity and genotyping accuracy when run on a single individual in isolation.

Taken together, the above analyses demonstrate that SpeedSeq achieves clinically relevant SV detection performance with excellent sensitivity – at least for SVs detectable with current technologies – clinically acceptable levels of accuracy, and highly accurate SV genotype information.

### Interpreting genomic variants

We have engineered SpeedSeq to seamlessly interface with GEMINI, a flexible software package for genome mining[21]. GEMINI is a local database framework that integrates variant calls (in VCF format) with information from various external databases including dbSNP, ENCODE, UCSC, ClinVar, and KEGG, allowing users to efficiently annotate and filter variants through succinct command line queries or a graphical browser interface. Published features of GEMINI enable users to filter variants based on predicted severity, and refine their significance with comparisons to ClinVar for health relationships or KEGG and HPRD catalogs for pathway burden enrichment. In concert with SpeedSeq, we have made numerous enhancements to GEMINI, particularly in handling structural variants and interpreting somatic mutations. Two new databases have been added to GEMINI: the COSMIC catalogue of somatic mutations in cancer[31] and DGIdb, the Drug-Gene Interaction database[33]. Users can interrogate these databases with several novel GEMINI operations designed to rapidly prioritize variants in a tumor-normal pair. The “gemini set somatic” command uses tunable stringency settings to classify somatic variants. These variants may be subsequently refined using the “gemini actionable mutations” command, which retrieves mutations predicted to have medium or high severity consequences in genes in the COSMIC cancer census database, and automatically reports their interactions with available therapeutic agents in DGIdb. In addition, GEMINI can now parse standard VCF files to quickly identify structural variants that alter gene dosage or interrupt transcripts. We have woven SV functionality into somatic mutation annotation to create the “gemini fusions” module, which reports putative somatic gene fusions affecting COSMIC cancer genes. SpeedSeq automatically prepares VCF files of SNVs, indels, and SVs by annotating them with the Ensembl Variant Effect Predictor (VEP)[20] so that they may be directly loaded into GEMINI.

### A case study: rapid-turnaround clinical analysis and interpretation of cancer genomes

Finally, to test SpeedSeq’s performance on real cancer data, we obtained deep (50X tumor, 30X normal) WGS data from five tumor-normal pairs (three colorectal, one ovarian, and one breast cancer) with validated somatic mutations from The Cancer Genome Atlas (TCGA) and processed them with SpeedSeq using default parameters. SpeedSeq recalled 96.4% of the orthogonally validated mutations across all five datasets including 98.8% of validated mutations in COSMIC census genes that have been causally implicated in cancer (Table 3). These are excellent levels of sensitivity considering that these mutations were originally ascertained by deep exome sequencing. To provide an example of a typical cancer analysis workflow, we next focused on an invasive breast carcinoma from TCGA that carries a known gene fusion[32]. We performed somatic variant calling on the tumor-normal pair using the standard SpeedSeq workflow. With four concise commands and less than an hour of computation, we loaded the VCF file into GEMINI, filtered variant calls for high-confidence, clinically relevant somatic mutations, and predicted gene fusion events (Fig. 6). This procedure produced several clinically actionable mutations in the tumor (including a previously confirmed missense mutation in TP53), as well as the known gene fusion event (Fig. 6). These analyses demonstrate the ease with which clinically relevant somatic mutations – including both point mutations and genomic rearrangements – can be identified using the SpeedSeq framework.

**Table 3.**
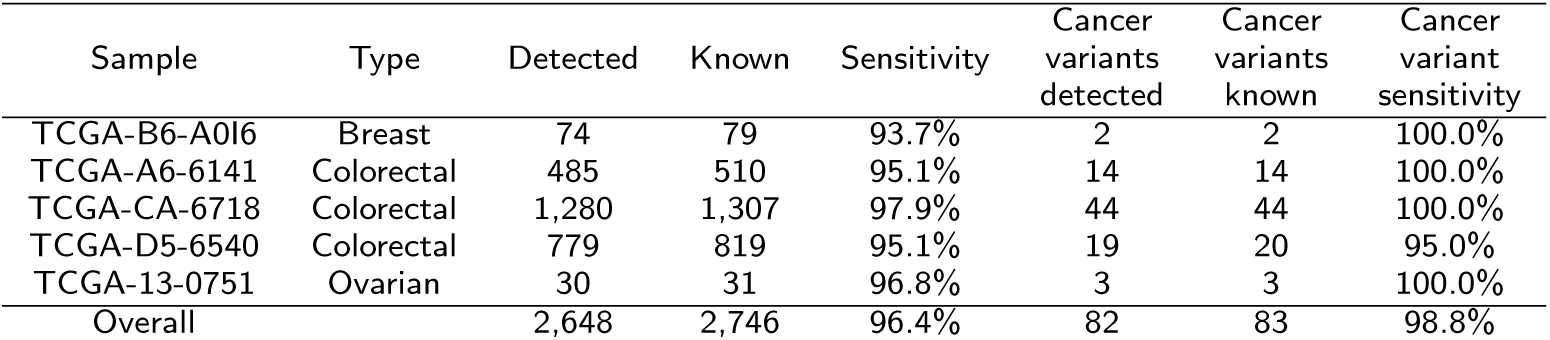
Sensitivity in detecting somatic mutations in tumor-normal pairs. We analyzed five tumor-normal pairs to assess SpeedSeq’s sensitivity in detecting 2,746 somatic mutations that had been previous detected by The Cancer Genome Atlas (TCGA) through deep exome sequencing and validated by an orthogonal method. The subset of cancer variants contains the variants within genes in the COSMIC cancer census gene set[31].

**Figure 6.**
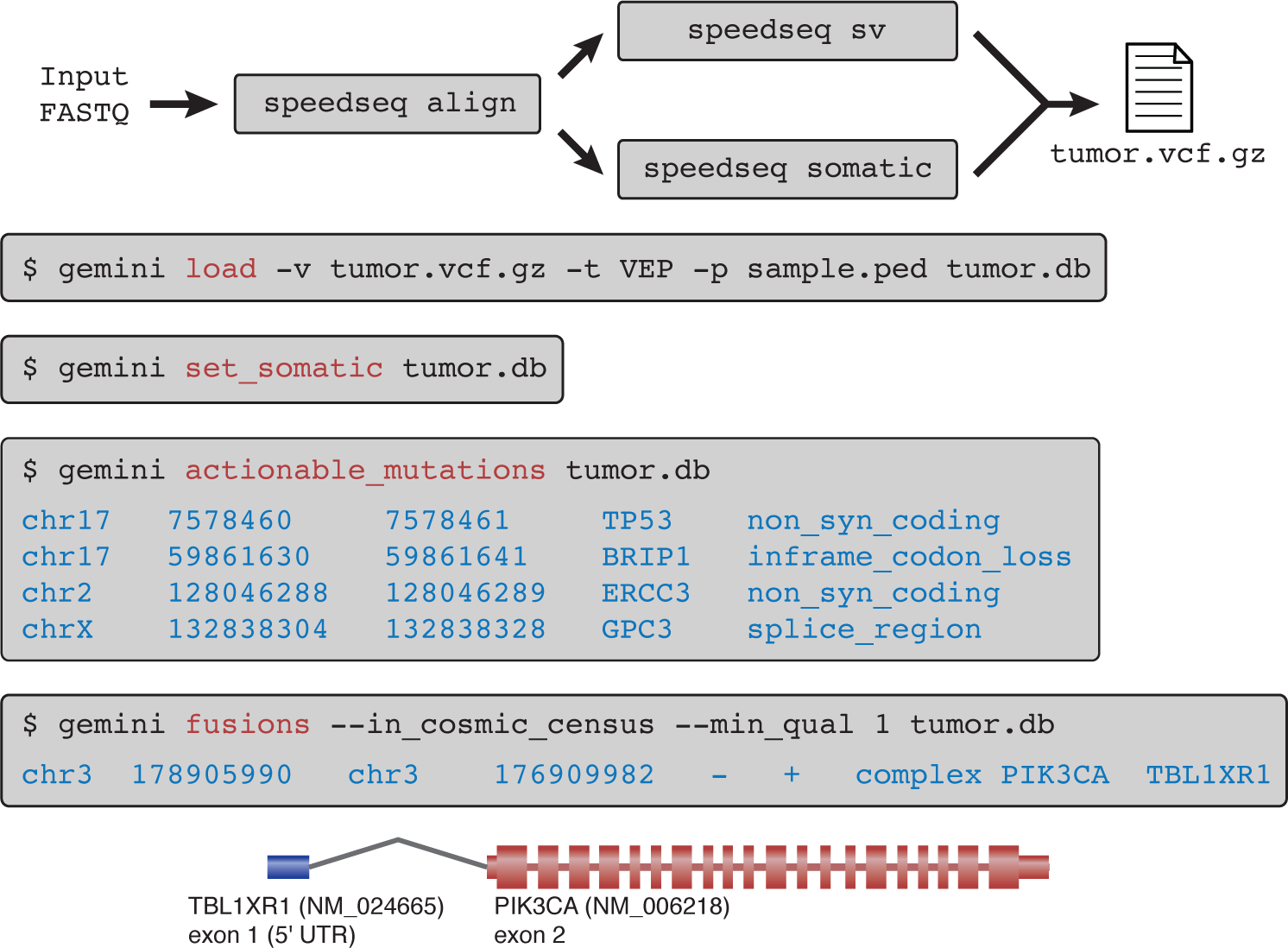
A case study in a tumor-normal pair. A SpeedSeq workflow demonstrating the seven succinct commands required to process a tumor-normal pair (TCGA-E2-A14P) from raw FASTQ reads to clinically actionable somatic mutations with predicted damaging consequences. In this tumor, SpeedSeq detected a somatic gene fusion product between exon 1 of TBL1XR1 and exon 2 of PIK3CA, a fusion that had been previously reported by Stransky *et al.*[32]

## Methods

### Alignment and BAM processing

SpeedSeq aligns paired-end FASTQ files to the human GRCh37 reference genome with BWA-MEM, using the -M flag to mark shorter alignments as secondary. In addition, several data manipulation components of typical pipelines are combined or reorganized to eliminate hours of processing. For example, duplicate marking, previously an 8 hour operation which prevents PCR amplification effects from biasing variant calling algorithms, is far more efficient when duplicates are marked on a query name-sorted BAM file streamed directly from the aligner. By streaming input directly from BWA, SAMBLASTER can seize idle CPU cycles that are periodically liberated each time BWA reads a FASTQ data chunk into the buffer. Furthermore, by marking duplicates at this earlier stage, this step can be combined with the extraction of discordant read-pairs and split read alignments, a process that also requires a query name-sorted BAM file. The SAMBLASTER tool achieves the fastest duplicate marking to date, while simultaneously producing auxiliary files for downstream analyses[15].

### SNV and indel calling

SpeedSeq runs FreeBayes version 0.9.16 with –min-repeat-entropy 1 and –experimental-gls parameters for germline variant calling. For somatic variant detection, SpeedSeq uses parameters tuned to increase sensitivity over low frequency variants (–pooled-discrete–genotype-qualities -F 0.05 -C 2 –min-repeat-entropy 1). SpeedSeq reports a somatic score (SSC) to estimate the confidence of each variant. The somatic score is the sum of the log odds ratios of the tumor (*LOD_T_*) and normal (*LOD_N_*) based on the genotype likelihood probabilities from FreeBayes.

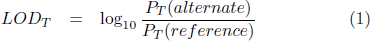

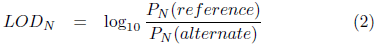

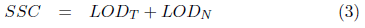

To increase speed, SpeedSeq runs a parallelized implementation of FreeBayes over 34,123 windowed regions of the genome averaging ∼84 kb in length. We generated these regions by partitioning the genome into bins of approximately equal numbers of reads, based upon the aggregate coverage depth of all 17 members of the CEPH 1463 family pedigree (Supplementary Fig. 1) (see Excluded regions). This binning scheme balances the computational load over the FreeBayes instances by allocating processors based on the quantity of expected input data. Additionally, it allows us to prioritize the processing hierarchy so that higher depth regions are analyzed first, further reducing the chance of unbalanced parallelization.

### Structural variation detection

SpeedSeq uses LUMPY to detect structural variant breakpoints from paired-end and split-read signals, and CNVnator[19] to detect copy number variants (CNVs) by read-depth analysis. Analyses presented here were performed using the SpeedSeq defaults (-mw 4 -tt 0, min_clip 20, min_non_overlap 101, min_mapping_threshold 20, discordant_z 5, back_distance 10, and weights of 1 for both paired-end and split-read evidence). SpeedSeq converts raw LUMPY output into a VCF file, merging evidence from multiple libraries by extracting the SM (sample) readgroup tag from each BAM file. This VCF file can be optionally annotated for copy number state at each structural variant with CNVnator version 3.0. Our custom parallelized implementation of CNVnator performs copy number segmentation on bins of 100 bp, and reduces processing time by allocating each chromosome to a separate parallel process.

### Structural variant genotyping

SVTyper is a maximum-likelihood Bayesian classification algorithm to infer an underlying genotype at each SV. Alignments at SV breakpoints either support the alternate allele with split or discordant reads, or they support the reference allele with concordant reads that bridge the breakpoint. The ratio and quantity of these observations allows probabilistic inference of genotype likelihood.

Under the assumption of diploidy, the set of possible genotypes at any locus is *G* = {*reference*, *heterozygous*, *homozygous*}. We defined the function *S*, where *S*(*g*) is the prior probability of observing a variant read in a single trial given a genotype *g* at any locus. These priors were set to 0.1, 0.4, 0.8 for reference, heterozygous, and homozygous deletions respectively. Assuming a random sampling of reads, the number of observed alternate (*A*) and reference (*R*) reads (scaled by mapping quality, 10^(−*mapq*/10))^ will follow a binomial distribution *B*(*A* + *R, S*(*g*^′^)), where *g*^′^ ∊ *G* is the true underlying genotype. Using Bayes’s theorem we can derive the conditional probability of each underlying genotype state from the observed read counts (Eq. 4). For these experiments, we have set the a priori probability 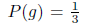 for each genotype, giving them equal expectations. Finally, we calculate 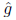 as the inferred genotype for the variant. Since the algorithm only interrogates SVs in the VCF file that have passed LUMPY filters as non-reference, it reports the more likely genotype of heterozygous or homozygous alternate states.

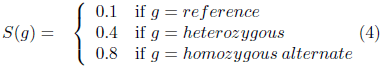

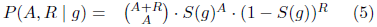

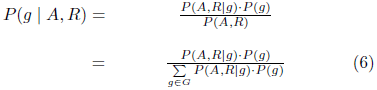

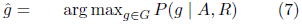

### Excluded regions

Despite the high quality of the human reference genome, artifacts remain in low-complexity regions and unannotated paralogous sequences that delay processing time and confound variant interpretation. These regions exhibit a several-fold increase in sequencing coverage depth where reads from disparate parts of the genome accumulate, violating the diploid assumption of downstream variant calling algorithms. To identify these high-depth regions we obtained PCR-free, 50X, paired-end sequencing data for all 17 members of the CEPH 1463 family pedigree in the Illumina Platinum Genomes project (European Nucleotide Archive: ERP001960). We aligned the reads to GRCh37 with BWA-MEM and measured aggregate coverage depth from all family members, excluding 15.6 Mb (0.6% of the genome) where the depth was greater than 2 × mode coverage + 3 standard deviations (Supplementary Fig. 1a). These criteria serve to exclude genomic regions that exhibit aberrantly high sequence coverage, while allowing for a 2-fold increase in copy number relative to the reference. These high-depth excluded regions are excluded from SNV, indel, and structural variant calling applications of SpeedSeq. For structural variation detection with LUMPY, we also exclude the mitochondrial genome, which is prone to false positive calls due to extremely high depth.

### SNV and indel validation

We compared SpeedSeq’s germline SNV and indel variant calling against two independent truth sets for NA12878, one derived from the Genome in a Bottle NA12878 gold standard calls and the other based on Omni microarray data from the 1000 Genomes Project. The GIAB 2.17 truth set contained 2,803,144 SNVs and 364,031 indels within highly confident regions (excluding segmental duplications, simple repeats, decoy sequence, and CNVs), spanning 2.2 Gb (77.6% of the mappable genome) for which non-variant sites could be confidently considered homozygous reference. The Omni microarray truth set contained 2,177,040 informative SNVs of which 689,788 were non-reference in NA12878, excluding markers within 50 bp of known indels. PCR-free 50X fastq files for NA12878 were obtained from the Illumina Platinum Genomes project were aligned and variant-called with SpeedSeq using to default parameters. To benchmark SpeedSeq’s performance against other standard tools, we also processed the aligned BAM files according to the Genome Analysis Toolkit (version 3.2-2-gec30cee) best practices workflow, including realignment around indels, base recalibration, and variant calling with Unified Genotyper (GATK-UG) and Haplotype Caller (GATK-HC). Variant quality score recalibration was performed on GATK variant callers, using a passing tranche filter of *<*99%. We normalized and compared variant calls according to the GIAB protocol, with vcfallelicprimatives, GATK’s LeftAlignAndTrimVariants, and VcfComparator[34]. We filtered variants for sensitivity and FDR against the GIAB truth set using a minimum quality score of 100 for GATK tools, and 1 for SpeedSeq (open circles, Fig. 2a,b). To evaluate performance in detecting somatic variants, we generated a simulated tumor-normal matched pair from the CEPH 1463 family Illumina Platinum data. The “tumor” dataset was an equal mixture of all 11 members of the F_2_ generation, downsampled to 50X coverage and aligned with SpeedSeq. The father of the F_2_ generation (NA12878) represented the 50X germline matched normal sample. Thus any variants present in the mother (NA12878) but absent from the father (NA12877) would appear as somatic mutations in the tumor-normal pair. In addition to retaining library preparation and chemistry biases of genuine sequence data, the heterozygous variants in the NA12878 follow a binomial pattern of inheritance in her children, allowing us to evaluate sensitivity and specificity for subclonal somatic mutations at a range of variant allele frequencies (Fig. 3a). To determine the expected allele frequency of somatic variants in the simulated tumor dataset, we selected variants that were present in the NA12878 GIAB 2.17 truth set but absent from NA12877 based on Real Time Genomics (RTG) joint variant callset of the entire CEPH 1463 pedigree (ftp://ftp-trace.ncbi.nih.gov/giab/ftp/data/NA12878/variant_calls/RTG/cohort-illumina-wgs.vcf.gz). We assessed the presence/absence of these variants in the F_2_ generation using RTG calls to derive their expected variant allele frequency in the simulated tumor dataset. For inclusion in the somatic SNV truth set, we required a variant to be diallelic, autosomal, and called by RTG as non-reference in NA12878 and reference in NA12877. Additionally, variants were disqualified from the truth set if they were called by RTG as homozygous for the alternate allele in any of the F_2_ children (a Mendelian violation). These criteria resulted in a set of 875,206 high confidence SNVs covering 77.6% of the mappable genome. We processed the data with SpeedSeq, MuTect 1.1.4, SomaticSniper, and VarScan 2 using parameters designed to target variants as low as 5% variant allele fraction. Receiver operating characteristic (ROC) curves were generated by varying somatic score (SSC) for SpeedSeq, SomaticSniper, and VarScan 2. For MuTect, which does not produce a single quality score for somatic variants, we varied the t_lod_fstar value to construct the ROC curve.

### Structural variant evaluation

We used inheritance information from the CEPH 1463 pedigree to evaluate deletion calls from SpeedSeq. Genuine deletions in the F_2_ generation are expected to segregate along SNV-based haplotype phasing of the chromosomes through all three generations of the pedigree, while variants that segregate discordantly can be assessed as false positives. To construct haplotype maps of the F_2_ genomes, we called SNVs with SpeedSeq on the entire 17-member pedigree, and phased SNVs by transmission at polymorphic sites in the parents. We smoothed the chromosomes for contiguous blocks of inheritance by selecting informative bases where 95% of each run of 101 SNVs reported a consistent parent-of-origin. We then merged regions that shared inheritance and were within 100 kb of each other. This allowed us to trace an average of 1.8 Gb (63.4%) of each F_2_ chromosome back to a particular grandparent, encapsulating meiotic crossovers that occurred in the F_1_ germline (Fig. 3). We then used SpeedSeq to jointly call structural variants on the entire pedigree, filtering for deletions that had at least seven pieces of support in at least one member of the pedigree, had legal Mendelian transmission, and whose origin could be unambiguously attributed to a single grandparent. Variants for which the founding grandparent by SV transmission agreed with the founding grandparent by SNV phasing were considered to be concordant, with strong supporting evidence for their authenticity.

### Tumor-normal pairs

Whole-genome sequencing data from five matched tumor-normal pairs and their orthogonally validated somatic mutations were obtained from The Cancer Genome Atlas (TCGA). These included three colorectal (TCGA-A6-6141, TCGA-CA-6718, TCGA-D5-6540), one ovarian (TCGA-13-0751), and one breast cancer (TCGA-B6-A0I6). Raw FASTQ reads were downsampled to 50X coverage in the tumor and 30X coverage in the normal sample. Samples were processed with SpeedSeq for alignment, somatic mutations, and structural variants using default parameters and then loaded into GEMINI for variant interpretation. We also analyzed whole genome sequencing data from a tumor-normal pair (∼63X tumor, ∼49X normal coverage) of a patient with an invasive breast carcinoma (TCGA-E2-A14P) containing a previously reported gene fusion between TBL1XR1 and PIK3CA[32].

### Hardware

All timings reported herein were performed on a single machine with 128 GB RAM and two Intel Xeon E5-2670 processors, each with 16 threads.

